# Mitochondrial CISD1/Cisd accumulation blocks mitophagy and genetic or pharmacological inhibition rescues neurodegenerative phenotypes in *Pink1/parkin* models

**DOI:** 10.1101/2023.05.14.540700

**Authors:** Aitor Martinez, Alvaro Sanchez-Martinez, Jake T. Pickering, Madeleine J. Twyning, Ana Terriente-Felix, Po-Lin Chen, Chun-Hong Chen, Alexander J. Whitworth

## Abstract

**Background:** Mitochondrial dysfunction and toxic protein aggregates have been shown to be key features in the pathogenesis of neurodegenerative diseases, such as Parkinson’s disease (PD). Functional analysis of genes linked to PD have revealed that the E3 ligase Parkin and the mitochondrial kinase PINK1 are important factors for mitochondrial quality control. PINK1 phosphorylates and activates Parkin, which in turn ubiquitinates mitochondrial proteins priming them and the mitochondrion itself for degradation. However, it is unclear whether dysregulated mitochondrial degradation or the toxic build-up of certain Parkin ubiquitin substrates is the driving pathophysiological mechanism leading to PD. The iron-sulphur cluster containing proteins CISD1 and CISD2 have been identified as major targets of Parkin in various proteomic studies.

**Methods:** We employed *in vivo Drosophila* and human cell culture models to study the role of CISD proteins in cell and tissue viability as well as aged-related neurodegeneration, specifically analysing aspects of mitophagy and autophagy using orthogonal assays.

**Results:** We discovered that the *Drosophila* homolog Cisd accumulates during aging, as well as in *Pink1* and *parkin* mutant flies. We observed that build-up of Cisd is particularly toxic in neurons, resulting in mitochondrial defects and Ser65-phospho-Ubiquitin accumulation. Age-related increase of Cisd blocks mitophagy and impairs autophagy flux. Importantly, reduction of Cisd levels upregulates mitophagy *in vitro* and *in vivo*, and ameliorates pathological phenotypes in locomotion, lifespan and neurodegeneration in *Pink1/parkin* mutant flies. In addition, we show that pharmacological inhibition of CISD1/2 by rosiglitazone and NL-1 induces mitophagy in human cells and rescues the defective phenotypes of *Pink1/parkin* mutants.

**Conclusion:** Altogether, our studies indicate that Cisd accumulation during aging and in *Pink1/parkin* mutants is a key driver of pathology by blocking mitophagy, and genetically and pharmacologically inhibiting CISD proteins may offer a potential target for therapeutic intervention.

## Background

Parkinson’s disease (PD) is a common neurodegenerative disorder caused by the progressive loss of midbrain dopaminergic (DA) neurons resulting in a spectrum of clinical symptoms including classic postural instability and loss of motor control as well as many other non-motor symptoms. The cause(s) of selective vulnerability of the neurons lost in PD is not precisely known, with genetic and epidemiological evidence primarily indicating defects in proteostasis, redox balance, calcium handling and mitochondrial homeostasis [1]. Compelling insights have come from understanding the pathogenic mechanisms of inherited forms of PD.

Mutations in the mitochondrial-targeted kinase PINK1 and the E3 ubiquitin ligase Parkin cause familial forms of parkinsonism [2,3]. A substantial body of evidence supports that PINK1 and Parkin act in a common pathway to promote the degradation of dysfunctional mitochondria through selective autophagy, termed mitophagy. This mechanism involves the stabilisation of PINK1 on the outer mitochondrial membrane (OMM) of dysfunctional mitochondria where it phosphorylates latent ubiquitin bound to OMM-resident proteins. This Ser65-phospho-ubiquitin (pUb) serves as a recruitment signal for Parkin, which is also phosphorylated by PINK1 at Ser65 to activate it, whereupon it additionally ubiquitinates OMM proteins for further phosphorylation triggering proteasomal degradation of specific targets and the recruitment of autophagy receptors, such as p62 [4–6]. In general, mitophagy is considered a key element of mitochondrial quality control (MQC) acting to safeguard a healthy mitochondrial network and prevent age-related neurodegenerative diseases, such as PD [7]. While many of the molecular details of PINK1/Parkin-mediated mitophagy have been elucidated in cell culture models, *in vivo* studies have presented a complex picture of the impact of PINK1/Parkin on physiological mitophagy [8–12]. Moreover, the reasons why loss of PINK1 and Parkin activity causes DA neurodegeneration are still not clear, preventing the rational design of targeted therapeutic interventions.

Genetic studies in *Drosophila* have delivered fundamental insights into the biology of *Pink1* and *parkin* and the *in vivo* consequences of their loss. Genetic *Pink1/parkin* null mutants present robust PD-related phenotypes, e.g. locomotor deficits and progressive DA neuron loss, resulting from profound mitochondrial disruption, most notably in flight muscle [3–5]. An unbiased proteomic screen for putative parkin substrates in *Drosophila* neurons highlighted the sole homologue of CISD proteins (here, termed Cisd) as a prevalent ubiquitination target [16]. Indeed, CISD1 and CISD2 have been reported as top Parkin substrates in several human cell culture studies [17–19], cementing these as *bona fide* targets. Moreover, CISD1 has also been found to be a key substrate of the mitochondrial deubiquitinase USP30 [20–23], which has been shown to counteract Parkin function during mitophagy [24]. However, their functional contribution to mitophagy or the downstream phenotypes of *Pink1/park* loss have not been studied. Therefore, we sought to study the role of the *Drosophila* Cisd and its human homologues in mitophagy and Pink1/parkin-related neurodegeneration.

CISD proteins (CISD1, CISD2 and CISD3 in vertebrates) belong to the NEET protein family, which are characterised by a unique iron-sulphur [2Fe-2S] cluster-coordinating CDGSH domain. This positions CISD proteins as important regulators of redox, electron and [2Fe-2S] cluster transfer reactions, vital for iron metabolism, mitochondrial respiration and the regulation of reactive oxygen species (ROS) production [25]. CISD1/mitoNEET and CISD2/NAF-1 contain a transmembrane domain by which they anchor to OMM and ER, respectively, forming homodimers in each organelle [26]. CISD3/MiNT contains two CDGSH domains acting as a monomer and localises to mitochondrial matrix [27]. CISD1 has been shown to be important for mitochondrial respiration, iron homeostasis and redox balance, as well as a key factor in the export of [2Fe-2S] clusters from mitochondria [28–31]. CISD2 is a causal gene for Wolfram syndrome, a severe neurodegenerative disease characterised by optic atrophy, deafness, dementia, and juvenile-onset insulin-dependent diabetes mellitus [32]. Furthermore, CISD2 has been shown to inhibit autophagy and regulate Ca^2+^ release from ER through direct interaction with IP3R, as well as perturbing mitochondrial respiration and cristae structure [33–35].

In this study, we report that *Drosophila* Cisd (also known as Dosmit) accumulates during aging and in *Pink1/parkin* mutant flies. Cisd overexpression is sufficient to drastically affect mitochondrial morphology, motor behaviour and lifespan, and is particularly toxic when accumulating in neurons. Mechanistically, Cisd accumulation blocks mitophagy and inhibits autophagy flux. Conversely, loss of *Drosophila* Cisd or human CISD1 or 2 is sufficient to promote mitophagy *in vivo* and *in vitro*. Consequently, loss of Cisd significantly rescues PD-related phenotypes in *Pink1/parkin* mutants, including motor function and DA neurodegeneration. Finally, we show that the small-molecule CISD inhibitors, rosiglitazone and NL-1, induce mitophagy in cells and are able partially rescue *Pink1/parkin* fly phenotypes. Thus, pharmacological targeting of CISD proteins represent a new avenue to boost mitophagy and potentially treat PD or other age-related neurodegenerative diseases.

## Materials and methods

### *Drosophila* stocks and husbandry

*Drosophila* were raised under standard conditions in a temperature-controlled incubator with a 12h:12h light:dark cycle at 25 °C and 65 % relative humidity, on food consisting of agar, cornmeal, molasses, propionic acid and yeast.

*Cisd KO* (*Cisd^−/−^*) and *UAS-Cisd* were kindly provided by Chun-Hong [36]. *UAS-ref(2)P* (p62) was generously provided by David Walker [37], *park^25^* mutants [13], and *UAS-mito-QC* [8] lines have been previously described. *Pink1^B9^* mutants [15] were kindly provided by J. Chung (Seoul National University). The following strains were obtained from Bloomington *Drosophila* Stock Center (BDSC, RRID:SCR_006457): *w*^1118^ (BDSC_6326), *da-GAL4* (BDSC_55850)*, nSyb-GAL4 (*BDSC_51635), *Mef2*-GAL4 (BDSC_27390), *TH*-GAL4 (BDSC_8848), *UAS-lacZ* (BDSC_8529), *UAS-hCISD1* (BDSC_77990), *UAS-hCISD2* (BDSC_76845), *UAS-Atg5 RNAi* (BDSC_27551), *UAS-Atg8a* (BDSC_84981), *UAS-GFP-mCherry-Atg8a* (BDSC_37749) and *UAS-mCherry-Atg8a* (BDSC_37750, [38]). The following strains were obtained from Vienna Drosophila Research Centre (VDRC): *UAS-LacZ RNAi* GD (v51446), *Cisd RNAi* GD (v33925) and KK (v104501).

### Antibodies and reagents

The following mouse antibodies were used: ATP5A (Abcam, ab14748, 1:10000), Ubiquitin (clone P4D1, Cell Signalling Technology, 3936, 1:1000), CISD2 (Proteintech, 66082-1-Ig, 1:1000), TOM20 (clone F-10, Santa Cruz, sc-17764, 1:1000), HA (Cell Signaling, 2367, 1:1000), tyrosine hydroxylase (Immunostar, 22491), Tubulin (Sigma, T9026, 1:5000). The following rabbit antibodies were used in this study: pS65-Ub (Cell Signalling Technologies, 62802S, 1:1000), GABARAP/Atg8a (Abcam, ab109364, 1:1000), ref(2)P/p62 (Abcam, ab178440, 1:1000), PINK1 (Cell Signaling Technology, 6946, 1:1000), CISD1 (Proteintech, 16006-1-AP, 1:1000), CISD2 (Proteintech, 13318-1-AP, 1:1000), FLAG (Cell Signaling, 2368S, 1:1000), Tubulin (Abcam, ab18251, 1:5000). Rat anti-Calnexin (clone W17077C, BioLegend, 699401).

The following secondary antibodies were used: Goat anti-mouse-HRP (Abcam, Ab6789-1, 1:10000), Goat anti-rabbit-HRP (Invitrogen, G21234, 1:10000), Goat anti-mouse-IRDye 680RD (Li-Cor, 926-68070), Goat anti-rabbit-IRDye 800CW (Li-Cor, 926-32211), Goat anti-mouse-AF405 (Invitrogen, A48255), Goat anti-mouse-AF488 (Invitrogen, A11001), Goat anti-mouse-AF594 (Invitrogen, A11005), Goat anti-rabbit-AF488 (Invitrogen, A11008) and Donkey anti-rat-AF488 (Invitrogen, A21208).

DMEM/F12, GlutaMAX Supplement (31331093, Gibco), DMEM, high glucose, GlutaMAX Supplement, pyruvate (31966047, Gibco), Fetal Bovine Serum (FBS) (10270106, Gibco), MEM Non-Essential Amino Acids Solution (100X) (11140050, Gibco), Penicillin-Streptomycin (10,000 U/mL) (15140122, Gibco), Opti-MEM (31985062, Gibco), Lipofectamine RNAiMAX (13778075, Invitrogen), Lipofectamine3000 (L3000008, Invitrogen), phosphate buffered saline (DPBS (1X), no calcium, no magnesium, 14190-169, Gibco), Formaldehyde (FA) (28908, Fisher Scientific Ltd) and Pierce BCA Protein Assay Kit-500 mL (23223, Life Technologies) were purchased from Thermo Fisher Scientific. The following reagents were obtained from Sigma: Dimethyl sulfoxide (DMSO) (276855-250ML), Rosiglitazone (R2408-50MG), Antimycin A (A8674-25MG). 5X siRNA Buffer (B-002000-UB-100, Horizon Discovery), mitoNEET Inhibitor NL-1 (475825-10MG, Calbiochem), cOmplete protease inhibitors (4693159001, Roche), PhosSTOP phosphatase inhibitors (4906837001, Roche), Bovine Serum Albumin (BSA, Fraction V, Cold-ethanol Precipitated, BP1605100, Fisher BioReagents), Oligomycin A (495455-10MG, Merck Life Science Limited), Chloroquine (C6628-25G, Merck Life Science Limited).

### Cell culture, transfection, RNA interference and drug treatment

ARPE-19 cells, stably expressing mCherry-GFP-Fis1(101–152) referred to as ARPE-19-MQC, were a kind gift of Dr. Ian Ganley (University of Dundee), and hTERT-RPE1, hTERT-RPE1-YFP-PARKIN cells were generously donated by Prof. Jon D Lane (University of Bristol). Cells were cultured in Dulbecco’s modified Eagle’s medium DMEM/F12 (hTERT-RPE1, hTERT-RPE1-YFP-PARKIN, and ARPE-19-MQC) or DMEM (U2OS) supplemented with 10% FBS, 1% non-essential amino acids and 1% penicillin/streptomycin. For siRNA experiments, cells were reverse-transfected with 20 nM of non-targeting (NT1) or target-specific siRNA oligonucleotides (Dharmacon On-Target Plus or siGenome, ThermoFisher Scientific), using Lipofectamine RNAi-MAX (Invitrogen) according to manufacturer’s instructions for 72h. Alternatively, cells were seeded and transfected for 48h with pcDNA3.1-CISD1-FLAG or pcDNA3.1-CISD2-FLAG using Lipofectamine3000 (Invitrogen) as per manufacturer instructions. For drugs treatment experiments, cells were treated with 100 µM of rosiglitazone or NL-1 for 24 h.

### siRNA and plasmids

siRNAs used in this manuscript were as follows: ONTARGETplus Non-Targeting siRNA oligo #1 (D-001810-01-20, Horizon Discovery), ON-TARGETplus Human CISD1 siRNA SMARTpool (L-020954-01-0020, Horizon Discovery), ON-TARGETplus Human CISD2 siRNA SMARTpool (L-032593-02-0020, Horizon Discovery). ORFs from hCISD1 and hCISD2 were amplified from transgenic flies carrying hCISD1 (BDSC_77990) or hCISD2 (BDSC_76845). FLAG was introduced at the C-terminal during the PCR amplification. Gibson assembly was used to introduce these PCR products into a pcDNA3.1 cut with KpnI and NotI. The following primers were used for the PCR amplification: hCISD1_KpnI_pCDNA31_Fwd: CTGGCTAGCGTTTAAACTTAAGCTTGGTACCATGAGTCTGACTTCCAGTTCCA hCISD1_FLAG_NotI_Rev: AAACGGGCCCTCTAGACTCGAGCGGCCGCCTACTTGTCATCGTCGTCCTTGTAGTCAGTTTCTTTTTTCTTGATGATCAGAG hCISD2_KpnI_pCDNA31_Fwd: CTGGCTAGCGTTTAAACTTAAGCTTGGTACCATGGTGCTGGAGAGCGTGG hCISD2_FLAG_NotI_Rev: AAACGGGCCCTCTAGACTCGAGCGGCCGCCTACTTGTCATCGTCGTCCTTGTAGTCTACTTCTTTCTTCTTCAGTATTAGTG.

### *Drosophila* and cell cultures harvesting and lysis

For the analysis of cell culture lysates by immunoblot, cells were rinsed once with cold PBS and subsequently lysed with 100 -200 μL cold RIPA buffer (50 mM Tris pH8, 180 mM NaCl, 1mM EDTA, 1% Triton-X100, 0.5% SDS), supplemented with cOmplete protease inhibitors and phosphatase inhibitors. Cell lysates were incubated on ice for 15-20min before being harvested into an Eppendorf tube. In the case of *Drosophila* lysates, animals (5 to 20 per replicate) with the right genotype were selected under CO_2_ and flash-frozen in liquid N_2_, all direct comparing genotypes being harvested equally at the same time. 150-200 µL of ice-cold RIPA buffer was added to 2 mL tubes containing 1.4 mm ceramic beads (15555799, Fisherbrand) to which flies were added and lysed using a Minilys homogeniser (Bertin Instruments). Flies were shaken twice at maximum speed, 10 seconds followed by a brief incubation on ice, after which samples were shaken again at maximum speed, 10 seconds, making a total of three cycles. After the lysis, *Drosophila* samples were returned to ice for at least 40 minutes. Both cell culture and *Drosophila* lysates were then centrifuged 5 minutes at maximum speed (21,000 x g) at 4 °C. Supernatant was then transferred to a fresh Eppendorf tube and centrifuged. This step was repeated until samples were completely cleared.

### Immunoblotting

Protein content was determined by BCA assay as per manufacturer instructions. Samples were then diluted in Laemmli x4 Sample Buffer (1610747, Bio-Rad) and analysed by SDS-PAGE using Mini-PROTEAN TGX Gels 4-20% (4561093, Bio-Rad) or similar. Gels were transferred onto precast nitrocellulose membranes (1704158, BioRad) using the BioRad Transblot Turbo transfer system, and membranes were immediately washed in distilled water and stained with ponceau (P7170-1L, Sigma) Membranes were then blocked by incubation with 5% (w/v) semi-skimmed milk in PBS 1X containing 0.1% (v/v) Tween-20 (PBStw) for 1 hour at least. Membranes were then incubated in a shaker at 4 °C overnight with appropriate primary antibodies in 5%milk-PBStw. 3 washes of 10min with PBStw were performed to rinse the membranes and then were incubated for one hour in secondary antibodies made in 5%milk-PBStw. Membranes were then washed 3 times,10min each, in PBStw. Membranes were developed either with ECL reagent (RPN2232, Amersham) using the Amersham Imager 680 (GE Healthcare), or alternatively with the LICOR secondary antibodies and imaged in an Odyssey LICOR imaging system. Image analysis was performed using of FIJI (Image J).

### Immunohistochemistry and sample preparation in *Drosophila*

*Drosophila* brains were dissected from 30-day-old flies and immune-stained with anti-tyrosine hydroxylase as described previously [39]. Tyrosine hydroxylase-positive neurons were counted under blinded conditions. For immunostaining of adult flight muscles and larval brains, animals were dissected in PBS 1X and fixed in 4% FA/PBS for 30 min at room temperature, permeabilized in 0.3% Triton X-100 for 30 min, and blocked with 1% BSA in 0.3% Triton X-100 PBS for 1h at RT. Tissues were then incubated in a shaker with the appropriate primary antibody: ATP5A antibody (Abcam, ab14748, 1:500), pUb (pS65-Ub, Cell Signalling Technologies, 62802S, 1:250) and/or ref(2)P/p62 (Abcam, ab178440, 1:1000), diluted in 1% BSA in 0.3% Triton X-100 PBS overnight at 4°C. Next day, samples were rinsed 3 times 10 min with 0.3% Triton X-100 in PBS, and incubated with the appropriate fluorescent secondary antibodies overnight at 4°C. The tissues were washed 3 times in 0.3% Triton X-100 in PBS followed by a last wash on PBS and mounted on slides using Prolong Diamond Antifade mounting medium (Thermo Fischer Scientific) and imaged next day. Tissues were imaged with a Zeiss LSM880 confocal microscope (Carl Zeiss MicroImaging) equipped with a Nikon Plan-Apochromat 63x/1.4 NA oil immersion objective. Images were analyses using FIJI (Image J).

### Mitochondrial morphology in larval brain

Third instar larvae overexpressing *UAS-mitoGFP* with the pan-neuronal driver *nSyb-GAL4* were dissected in PBS and fixed in 4% FA/PBS for 30 min at room temperature. The tissues were washed 3 times in PBS and mounted on slides using Prolong Diamond Antifade mounting medium (Thermo Fischer Scientific) and image next day in a Zeiss LSM880 confocal microscope (Carl Zeiss MicroImaging) equipped with Nikon Plan-Apochromat 63x/1.4 NA oil immersion objectives.

### Locomotor and lifespan assays

The repetitive iteration startle-induced negative geotaxis (RISING, or ‘climbing’) assay was performed using a counter-current apparatus. Experiments were performed using multiple small groups of flies of the indicated genotypes and ages. Briefly, flies were collected under minimal CO_2_ anaesthesia 24 hours before the assay. On the day of assay, flies are transferred to assay tubes and allowed to acclimatise in the behaviour room for ∼1h. Cohorts of ∼20 flies (25 max.) were placed into the first chamber of the counter-current apparatus, tapped to the bottom and given 10 s to climb a 10 cm distance. Flies that climbed >10 cm within this time were shifted to the neighbouring tube. This procedure was repeated five times, and the number of flies that remained in each chamber were counted. The weighted performance of several groups of flies for each genotype was normalised to the maximum possible score and expressed as the *climbing index* [13]. To age flies for climbing assays, they were transferred to fresh tubes every 2–3 days. For lifespan experiments, flies were grown under identical conditions at low density. Progeny were collected under very light (<30 s) anaesthesia and kept in tubes of 10–25, transferred to fresh media every 2–3 days and the number of dead flies recorded. Percent survival was calculated at the end of the experiment after correcting for any accidental loss or escape using https://flies.shinyapps.io/Rflies/ (Luis Garcia).

### Image analysis and quantification of mitolysosomes in *Drosophila* tissues

For mitolysosome imaging in *UAS-mito-QC*, tissues were dissected in PBS, fixed in 4% FA/PBS at pH 7 for 30min at room temperature and mounted. Third instar larval and adult brains were mounted in Prolong Diamond Antifade mounting medium (Thermo Fischer Scientific). While thoraces were mounted in VECTASHIELD Antifade Mounting Medium (H-1000, Vector Laboratories). Larval and adult brains were imaged using Andor Dragonfly spinning disk microscope (Oxford Instruments Group), equipped with a Nikon Plan-Apochromat 100x/1.45 NA oil immersion objective and iXon camera taking 10-13 µm z-stacks with 0.2 µm step size. Adult thoraces were imaged with a Zeiss LSM 880 confocal microscope (Carl Zeiss MicroImaging) equipped with Nikon Plan-Apochromat 63x/1.4 NA oil immersion objectives, taking 10 µm z-stacks with 1 µm step size. We find that the time elapsed from dissection-mounting and imaging is critical for the retrieval of red-only dots. Tissue specific optimisation was required. Larval brains were imaged early next day after dissection, while adult brains and thoraces were imaged briefly after (2-4 h later). Image analysis of larval brains was done employing Imaris (version 9.7.0) analysis software (BitPlane; RRID:SCR_007370) as previously described [8]. For adult *Drosophila* brains and thoraces, as well as for ARPE-19-MQC cells, mitolysosomes were analysed using the FIJI plug-in *mito-QC Counter* [40] as advised by authors. Analysis was performed in maximum projection images of *Drosophila* adult brain and ARPE-19-MQC cells, while *Drosophila* adult thoraces were analysed as per single plain.

The average number of mitolysosomes per cell was analysed per animal in larval and adult brain, as well as in ARPE-19-MQC cells, while mitolysosomes per area (µm) was calculated in *Drosophila* muscles. Data points in the quantification charts show the average mitolysosomes per biological replicate, where *n* ≥ 6 animals or *n* ≥ 60 cells for each condition.

### Transmission electron microscopy

Thoraces were dissected in 0.1 M cacodylate buffer and fixed in 1% glutaraldehyde and 4% paraformaldehyde in 0.1 M cacodylate buffer overnight at 4 °C. After washing with 0.1 M cacodylate buffer, the samples were treated with 1% OsO_4_ for 1 h at 4 °C. After washing with ddH_2_O, the samples were dehydrated in a gradient series using 30%, 50%, 70%, 95%, and 100% ethanol solutions at room temperature. The samples were then infiltrated using series epoxy resin at room temperature and mounted in pure resin. After polymerization at 60 °C for 2 days, 90 nm sections were prepared and observed with a Tecnai G2 Spirit TWIN transmission electron microscope (FEI, Hillsboro, OR, USA).

### Imaging and analysis of autolysosomes using GFP-mCherry-Atg8a reporter

#### Drosophila

Adult muscle from GFP-mCherry-Atg8a reporter *Drosophila* were dissected in PBS, fixed with 4% FA/PBS at pH 7 for 30 minutes at room temperature, and mounted in Prolong Diamond Antifade mounting medium (Thermo Fischer Scientific). Samples were imaged the day after. Confocal images, acquired with a Zeiss LSM 880 microscope (Carl Zeiss MicroImaging) equipped with Nikon Plan-Apochromat 63x/1.4 NA oil immersion objectives, taking 10 µm z-stacks with 1 µm step size, were processed using FIJI plug-in *mito-QC Counter* [40]. The quantification of autolysosomes was performed in a plane-by-plane basis similarly to mitolysosome analysis in muscles. Data points in the quantification charts show average autolysosomes per area (µm) for individual animals, where *n* ≥ 6 animals for each condition.

#### ARPE-19-mito-QC cell based mitophagy analysis

ARPE-19-mito-QC-FIS1 (Ian Ganley, University of Dundee) cells were used to assess mitophagy in a human cell model. After 48h knockdown with siRNA oligos cells were seeded onto an ibidi dish (IB-81156, ibidi, Thistle Scientific Ltd), and at 72h post-knock down cells were imaged live using a spinning disk microscope. Generated images were processed using the FIJI plug-in *mito-QC Counter* [40] as previously described. For treatments with rosiglitazone and NL-1: cells were seeded onto an ibidi dish, next day were treated with 100 µL rosiglitazone or NL-1 for 24h and imaged and analysed as described above.

#### Mitochondrial enrichment by differential centrifugation

All steps were performed on ice or at 4 °C. 20-30 flies were prepared either fresh or after flash-freezing in liquid nitrogen, with all direct comparisons performed with flies that were prepared in the same manner. Flies were transferred into a Dounce homogeniser containing 400 μL of mitochondrial isolation buffer (225 mM mannitol, 75 mM sucrose, 5 mM HEPES, 0.1 mM EGTA, pH 7.4) containing cOmplete protease inhibitors (Roche) and PhosSTOP phosphatase inhibitors (Roche), and manually homogenised with 15 strokes of a pestle. The homogenate was transferred to an Eppendorf tube, after which further 400 μl of mitochondrial isolation buffer was added to the homogeniser and the sample was homogenised with a further 5 strokes. The homogenates were pooled and centrifuged at 1500 g at 4°C for 6 min before being filtered through a 70 μm nylon cell strainer (352350, Falcon). The sample was then centrifuged at 7000 g at 4°C for 6 min and the resulting mitochondrial pellet was washed with isolation buffer once and finally resuspended into 150-200 μL RIPA buffer. The protein content was determined by BCA assay (Thermo Pierce).

#### USP2 deubiqutination assay

For the assessment of ubiquitinated Cisd in mitochondrial fractions, USP2 deubiquitinating assay was performed as previously described [41]. Briefly, whole fly mitochondrial fractions. 30 μg protein per mitochondrial fraction was treated with the pan-specific deubiquitinase USP2 (E-506, BostonBiochem). The USP2 enzyme was diluted in buffer (50 mM Tris–pH 7.5, 50 mM NaCl, 10 mM DTT) and then added to the subcellular fractions to a final USP2 concentration of 1 μM. The mixture was incubated for 45 min at 37°C prior to analysis by immunoblotting.

#### Rosiglitazone treatment in *Drosophila*

Parental crosses were set into food containing 1 mM rosiglitazone diluted 1:1000 or DMSO as control. The progeny of 0-3 days old flies was anaesthetised with CO_2_ and the correct genotype was selected for further investigation.

#### Light microscopy imaging of *Drosophila* thorax indentations and wing posture

0-3 days old flies were anaesthetised with CO_2_ and thorax indentations and abnormally extended wing posture were assessed by light microscopy imaging using a Nikon SMZ stereo zoom microscope fitted with 1x Apo lens. The number of animals displaying each phenotype was counted in a binary manner (present-absent) respective to the total.

#### Statistical analysis

Data are reported as mean ± SEM or mean ± 95% CI as indicated in figure legends. The n numbers of biological replicates are shown in the graphs or figure legend. For statistical analyses of lifespan experiments, a log rank (Mantel-Cox) test was used. For climbing assay analysis, a Kruskal-Wallis nonparametric test with Dunn’s post-hoc correction for multiple comparisons was used. Where multiple groups were compared, statistical significance was calculated by one-way ANOVA with Bonferroni post-test correction for multiple samples. When only two groups were compared a Welsch’s *t* test was used. All the samples were collected and processed simultaneously and therefore no randomization was appropriate. Unless otherwise indicated, images analysis was done in blind conditions. Statistical analyses were performed using GraphPad Prism 9 software (GraphPad Prism, RRID:SCR_002798). Statistical significance was represented in all cases as follows: *\* P* < 0.05, *\*\* P* < 0.01, *\*\*\* P* < 0.001 and **** *P* < 0.0001.

## Results

### *Drosophila* Cisd accumulates with age, disrupts mitochondria and is neurotoxic

Since Cisd and its homologues have been identified as parkin substrates, we hypothesised that Cisd abundance could be increased in *Drosophila parkin* mutants. Examining the steady-state levels of Cisd by immunoblotting we found Cisd levels were increased in aged *parkin* null (*park*^−/−^) and *Pink1* null mutants (*Pink1*^−/−^) (Fig. 1A, B). Mammalian CISD1 and CISD2 have been shown to form homodimers with high stringency [19,42]. We also observed Cisd dimers in our immunoblots even under standard reducing conditions (Fig. 1A). However, in non-reducing conditions, Cisd is almost exclusively found as a dimer (Fig. S1A). We confirmed that the upper band was indeed a dimerised form of Cisd and not a ubiquitinated form as mitochondrial fractions treated with the promiscuous deubiquitinase USP2 retain the dimer band (Fig. S1B). We also noted that Cisd levels increased with age in wild-type (WT) flies (Fig. 1A, C), as previously described [36], with both monomer and particularly the dimer increasing in abundance (Fig. S1A). Age-dependent increase in Cisd was particularly notable in fly head lysates (Fig. S1C-D).

**Figure 1.**
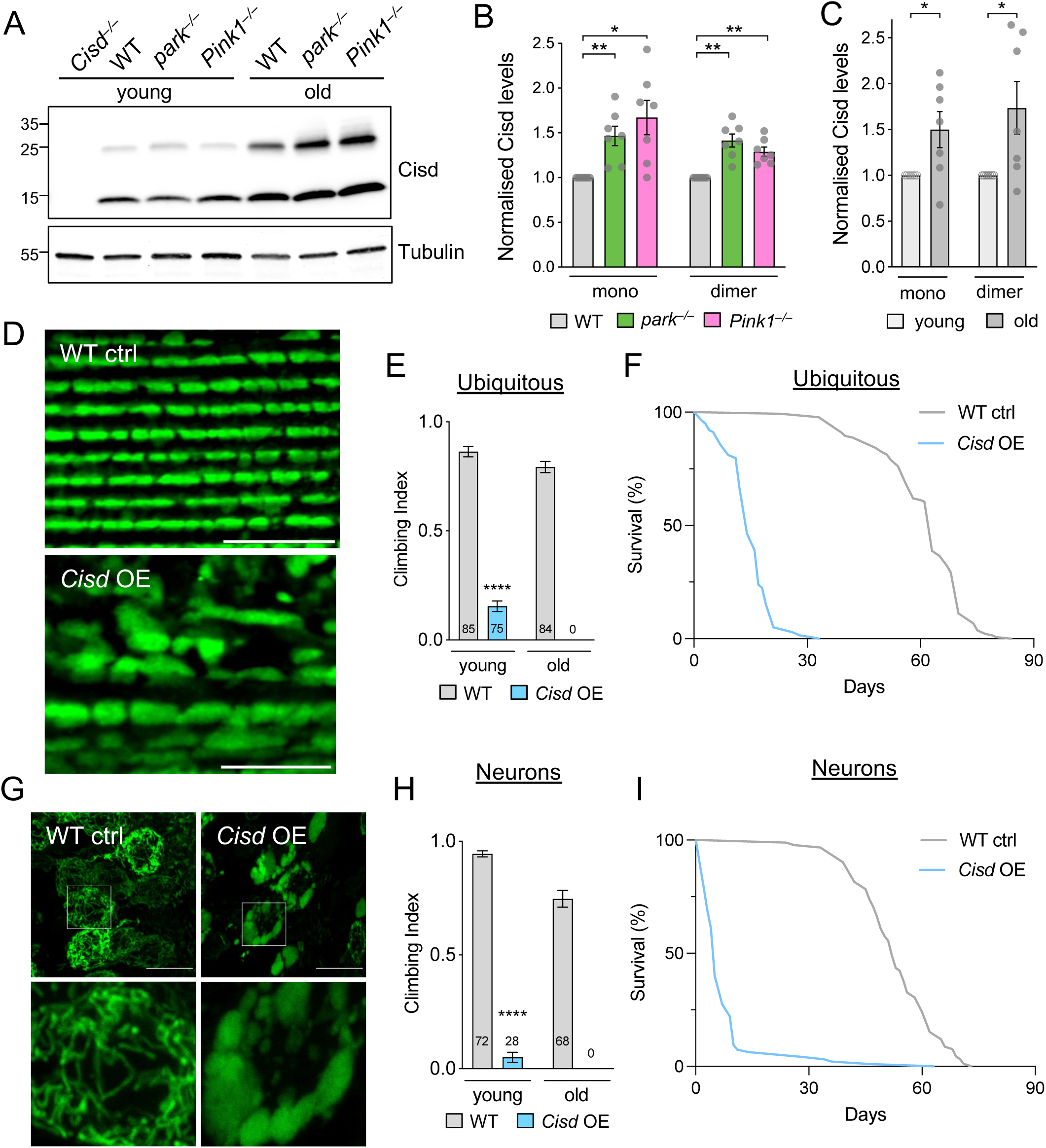
Cisd accumulation disrupts mitochondria affecting locomotion and lifespan. (A) Representative immunoblot of whole fly lysates of the indicated genotypes from young (2 days) and old (30 days) flies probed for Cisd and Tubulin as loading control. (B, C) Quantification of Cisd monomer or dimer levels analysed in A. (D) Confocal microscopy of flight muscle from (young) wild-type control (WT ctrl) or *Cisd* overexpressing (OE) flies immunostained for ATP5A mitochondrial marker. (E) Climbing assay of young or old WT and ubiquitous *Cisd* OE flies via the *da*-GAL4 driver. Charts show mean ± 95% CI. N is shown inside bars. (F) Lifespan of WT and ubiquitous Cisd OE flies. N > 130 animals. (G) Confocal microscopy of neuronal soma from control (WT ctrl) or *Cisd* overexpressing (OE) larvae co-expressing the mito.GFP mitochondrial marker via the *nSyb*-GAL4 driver. (H) Climbing assay of young or old WT and neuronal *Cisd* OE flies via the *nSyb*-GAL4 driver. Charts show mean ± 95% CI. N is shown inside bars. (I) Lifespan of WT and neuronal *Cisd* OE flies. N > 90 animals. Statistical analyses: (B) RM one-way ANOVA with Geisser-Greenhouse correction; (C) paired t-test; (E, H) Mann-Whitney non-parametric test. * *P* < 0.05; ** *P* < 0.01; **** *P* < 0.0001. Scale bars = 10 μm.

As Cisd levels increased with age, and the Pink1-parkin pathway acts to restrict its steady-state levels, we reasoned that elevated Cisd levels may be a key contributor to age-related organismal decline. Thus, we induced transgenic overexpression of *Cisd* (Fig. S1E), with different tissue-specific drivers, and assessed mitochondrial integrity and organismal fitness and survival. *Cisd o*verexpression via the moderate, ubiquitous driver *da-*GAL4 caused gross disruption to mitochondrial morphology, resulting in abnormally enlarged mitochondria in flight muscles (Fig. 1D), as previously reported [36]. It also severely impacted locomotor function and lifespan (Fig. 1E, F). Pan-neuronal overexpression of *Cisd* via *nSyb*-GAL4 also severely affected mitochondrial morphology and locomotion, and dramatically shortened lifespan (Fig. 1G-I). Notably, overexpression restricted just to DA expressing cells via *TH*-GAL4 also caused an age-related decline in motor ability (Fig. S1F), while muscle directed overexpression only caused a modest impact on locomotion and did not significantly affect lifespan (Fig. S1G, H).

Together, these data indicate that Cisd levels accumulate with normal ageing, as well as in *Pink1* and *parkin* mutants, and that Cisd accumulation (especially neuronal accumulation) is sufficient to substantially impact organismal vitality.

### *Drosophila* Cisd is functionally more related to mitochondria-localised CISD1

Both CISD1 and CISD2 have been reported as substrates of Parkin in proteomic studies [17–19]. Evaluating the dynamics of their degradation during PINK1/Parkin-mediated mitophagy, we treated retinal pigment epithelial (RPE1) cells (with endogenous Parkin) or cells overexpressing YFP-Parkin (RPE1-Parkin) with antimycin A and oligomycin to induce mitophagy and assessed CISD1 and 2 over time (Fig. 2A). Monitoring PINK1-mediated phospho-ubiquitin (pUb) showed rapid induction of mitophagy that steadily increased over time with a concomitant gradual decrease in the OMM protein TOM20 (Fig. 2A). As expected, pUb deposition and TOM20 loss was accelerated in Parkin-overexpressing cells. In these conditions, CISD1 was rapidly degraded during mitophagy similar to TOM20, however, we found that CISD2 levels were minimally affected (Fig. 2A). Co-localisation studies analysing the distribution of FLAG-tagged CISD1 or 2 (Fig. 2B) showed that while the majority of CISD1 is localised to mitochondria (TOM20 staining), CISD2 is predominantly localised to the ER (Calnexin staining). These observations are consistent with other reports [28,26] and align with the degradation of mitochondrial CISD1 while ER-localised CISD2 is largely spared during induced mitophagy.

**Figure 2.**
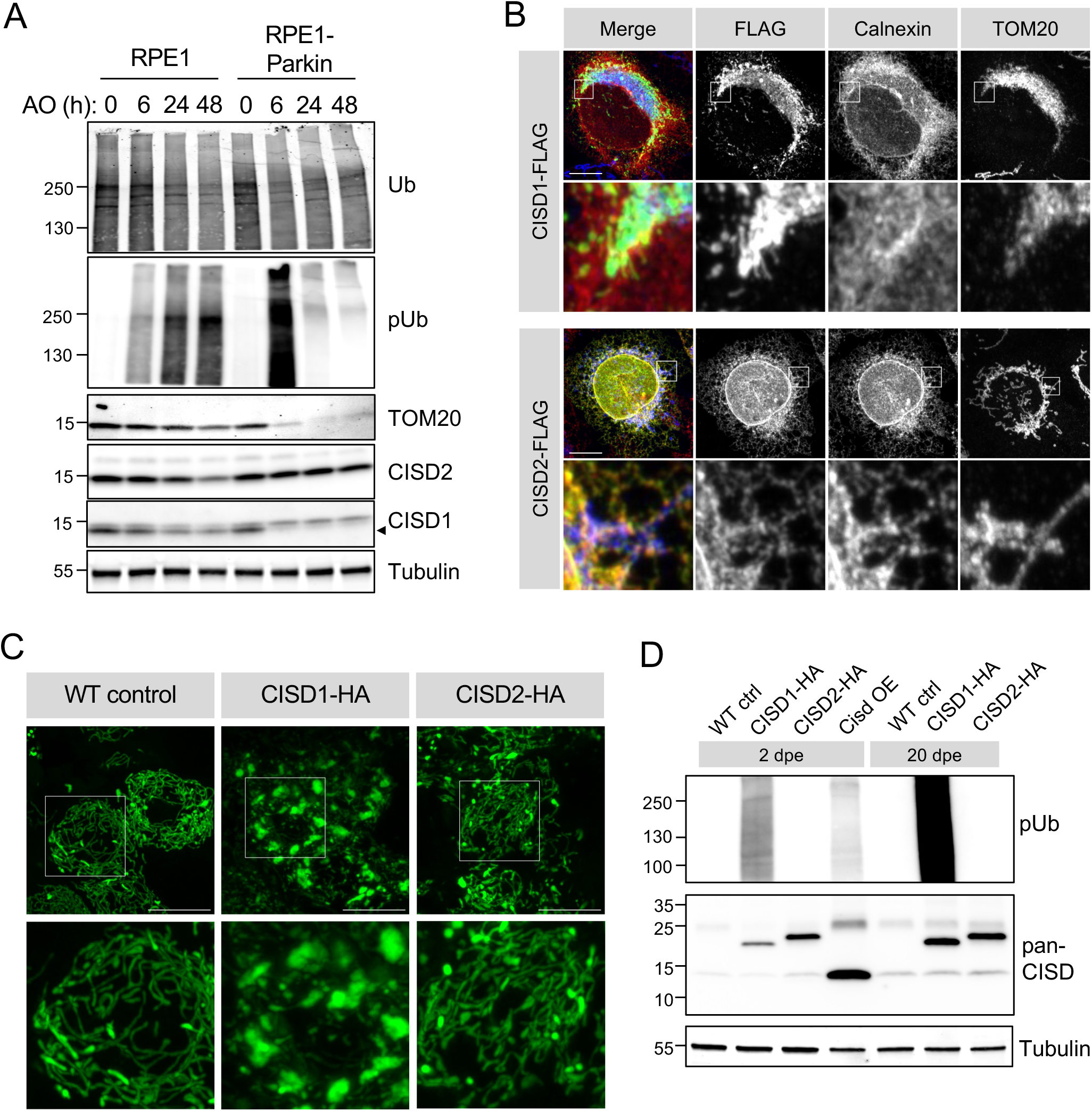
*Drosophila* Cisd is functionally more similar to CISD1. (A) Immunoblots of protein lysates from RPE1 cells ± YFP-Parkin overexpression treated with antimycin A (4 μM) and oligomycin (10 μM) for the indicated time to induce mitophagy, probed for mitophagy marker pUb, and degradation of TOM20 and CISD1/2, alongside respective loading control total Ub and Tubulin. Blot is representative of 3 replicate experiments. (B) Confocal micrographs of U2OS cells transfected with human CISD1-FLAG or CISD2-FLAG, counter-stained with antibodies against TOM20 (mitochondria) or Calnexin (ER). (C) Confocal micrographs of *Drosophila* larval neurons expressing transgenic mito.GFP with human CISD1-HA or CISD2-HA. (D) Immunoblots of protein lysates of whole flies expressing the indicated transgenes versus WT control, probed with the indicated antibodies. Scale bars = 10 μm.

*Drosophila* encode a single CISD orthologue with substantial amino acid sequence homology to human CISD1 and CISD2, with marginally higher homology to CISD2 than CISD1 (45% identity, 66% similarity vs CISD2; 42% identity, 65% similarity vs CISD1). Thus, we sought to determine whether *Drosophila* Cisd may function more similarly to one or other of the human orthologues. Ectopically expressing HA-tagged human *CISD1* and *CISD2* in *Drosophila* neurons, we found that expression of *CISD1* induced the same mitochondrial hyperfused phenotype (Fig. 2C) as seen with overexpression of *Cisd* (Fig. 1G), while *CISD2* expression had minimal effect (Fig. 2C). We previously noted that overexpression of *Drosophila Cisd* triggers mitochondrial pUb accumulation (discussed further below), this was mirrored by expression of *CISD1* but not *CISD2* (Fig. 2D). These results strongly support that *Drosophila* Cisd is functionally more related to CISD1 than CISD2.

#### *Cisd* overexpression prevents mitophagy and causes autophagosome accumulation

We recently described that Pink1-mediated pUb can be readily detected in *Drosophila* by immunoblotting or immunohistochemistry [41], and noted that pUb accumulates on dysfunctional mitochondria in *parkin* mutants (Fig. 3A, B). Here, we found that *Cisd* overexpression is also sufficient to cause a similar accumulation of pUb by immunoblotting (Fig. 3 A). Immunohistochemistry showed a concomitant accumulation of pUb surrounding selected mitochondria in flight muscle (Fig. 3B). The build-up of pUb is consistent with Cisd accumulation blocking efficient mitochondrial degradation and mirrors blocked mitophagy due to the loss of parkin. To verify these observations using an orthogonal approach we analysed mitophagy flux using the *mito*-QC mitophagy reporter flies [8]. This revealed that *Cisd* overexpression caused a significant reduction in mitolysosome number in larval and adult neurons (Fig. 3C-F). Thus, *Cisd* overexpression is sufficient to block mitophagy flux.

**Figure 3.**
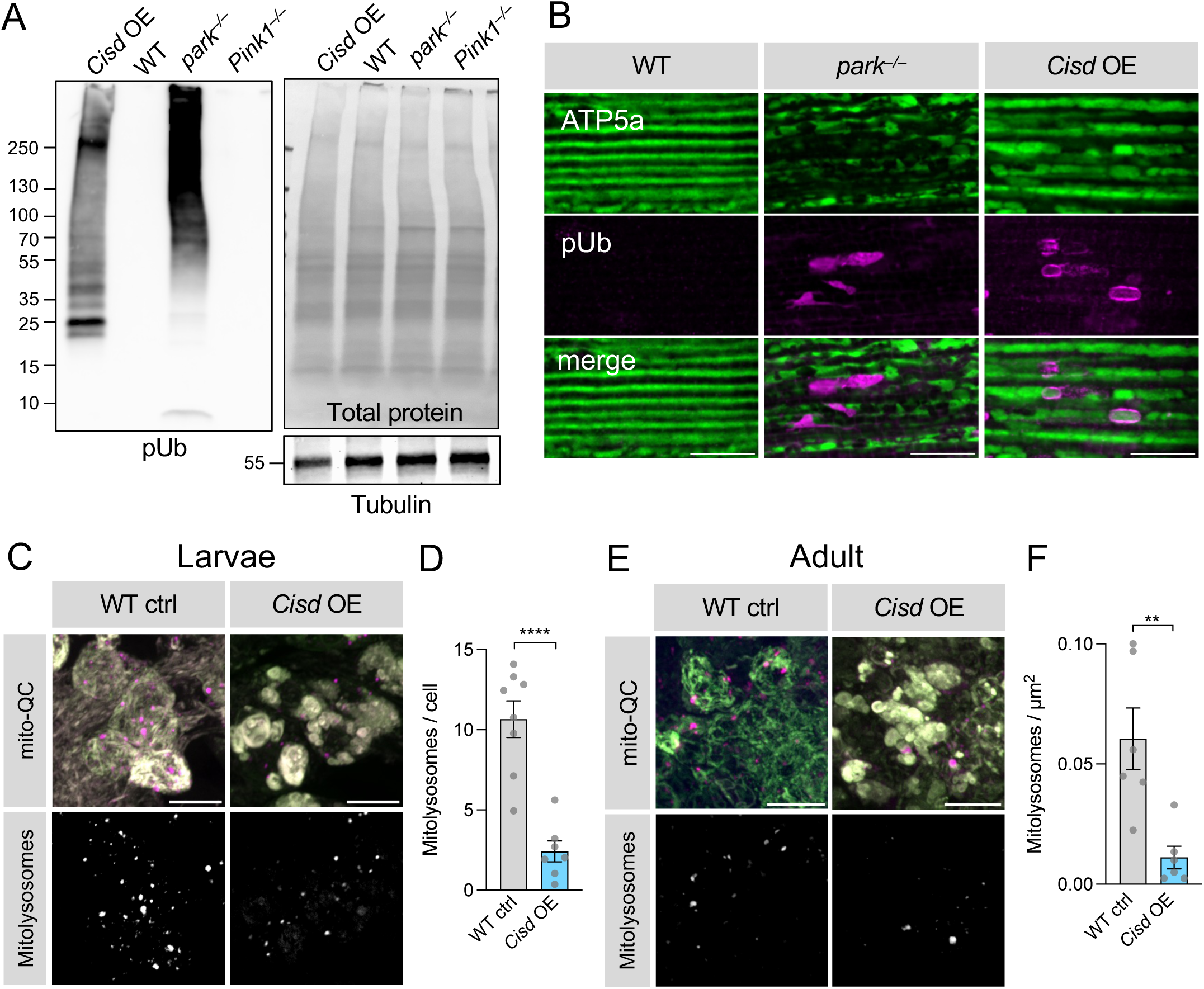
*Cisd* overexpression blocks mitophagy flux. (A) Immunoblot analysis of whole fly lysates from young flies of the indicated genotypes, analysed for pUb, and Tubulin or total protein levels as loading controls. Blot is representative of 3 replicate experiments. *Cisd* overexpression was driven by *da*-GAL4. (B) Confocal microscopy analysis of adult *Drosophila* flight muscle from young flies of the indicated genotype immunostained for mitochondria (ATP5A) or pUb. (C-F) Confocal analysis of mitophagy reporter *mito*-QC (OMM-localised tandem RFP-GFP) in larval (C, D) or adult (E, F) neurons of the indicated genotypes with ‘red-only’ mitolysosomes shown. (D, F) Number of mitolysosomes quantified shown in C and E. Data points indicated individual animals analysed. Statistical analysis: unpaired t-test; ** *P* < 0.01; **** *P* < 0.0001. Scale bars = 10 μm.

We postulated that the mitophagy block caused by Cisd accumulation may be due to defective autophagy. To investigate this we immunostained adult flight muscle overexpressing *Cisd* for the autophagy receptor p62 (also called ref(2)P in *Drosophila*) alongside the autophagosome reporter, mCherry-Atg8a (*Drosophila* homologue of LC3). This showed that p62-and Atg8a-positive puncta accumulate close to mitochondria, forming large aggregates (Fig. 4A). Analysing these structures by electron microscopy confirmed that indeed numerous autophagosomes accumulate around abnormally enlarged and vesiculated mitochondria in *Cisd* overexpressing tissue (Fig. 4B).

**Figure 4.**
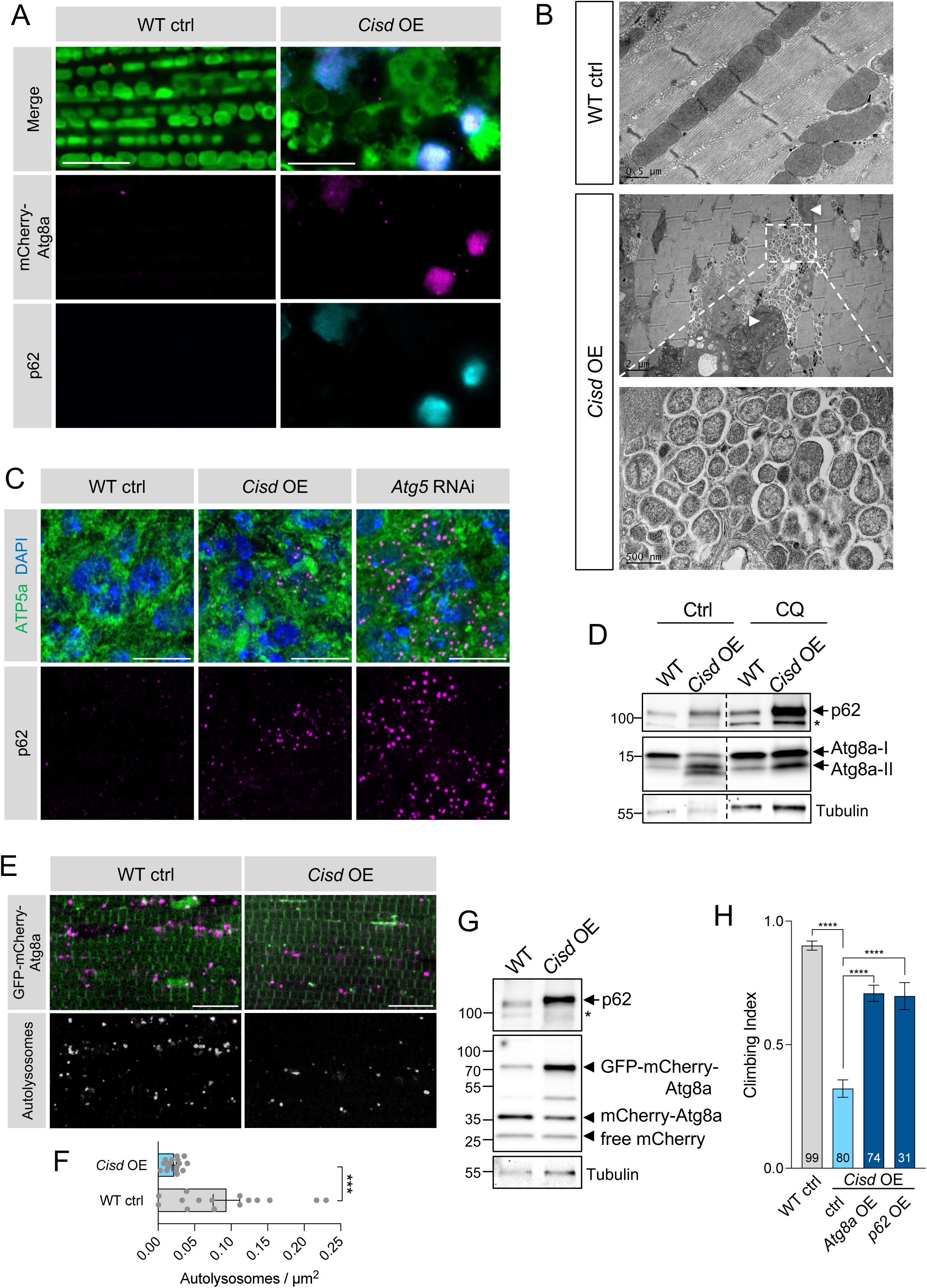
*Cisd* overexpression causes autophagosome accumulation and prevents autophagy. (A) Confocal micrographs of WT control versus *Cisd* overexpressing adult flight muscle, immunostained for p62 alongside imaging mCherry-Atg8a autophagosome reporter. (B) Electron micrographs of flight muscle as in A, showing multiple autophagic vesicles (inset) in proximity to disrupted mitochondria (arrowheads). (C) Larval neurons of WT control versus *Cisd* overexpressing or *Atg5* knockdown (via *nSyb*-GAL4 driver) animals immunostained p62 alongside ATP5a (mitochondria) and DAPI. (D) Immunoblot analysis of protein lysates from whole flies overexpressing *Cisd* or WT controls under basal conditions (Ctrl) or treated with 10mM chloroquine (CQ) for 24h. Blots were probed with antibodies against p62, Atg8a (LC3) and Tubulin. (E) Confocal microscopy analysis of adult flight muscle WT control versus *Cisd* overexpressing animals co-expressing the autophagy flux reporter GFP-mCherry-Atg8a. (F) Quantification of the number of autolysosomes shown in shown in E. Data points indicated individual animals analysed. Statistical analysis: unpaired t-test; *** *P* < 0.001. (G) Immunoblot analysis of protein lysates from whole flies expressing the autophagy reporter GFP-mCherry-Atg8a in the presence of WT control (WT) or *Cisd* overexpression. Blots were probed with antibodies against p62, mCherry and Tubulin. (H) Locomotor climbing assay of 2-day-old adult flies of the indicated genotypes. Charts show mean ± 95% CI. N is shown inside bars. Statistical analyses: Kruskal-Wallis non-parametric test with Dunn’s post-hoc correction. **** *P* < 0.0001. Scale bars = 10 μm for light microscopy, or indicated on image for EM.

Immunostaining larval brains also revealed an accumulation of p62 puncta upon *Cisd* overexpression, similar to that seen in autophagy-deficient *Atg5* RNAi flies (Fig. 4C). Likewise, immunoblot analysis of *Cisd* overexpressing flies showed increased levels of p62 and accumulation of the lipidated form of *Drosophila* LC3 (Atg8a-II), that didn’t further increase in the presence of the lysosomal inhibitor chloroquine (Fig. 4D), indicative of blockage in autophagy flux.

Consistent with these observations, the autophagy flux reporter, GFP-mCherry-Atg8a, also showed that autolysosome number was significantly decreased in *Cisd* overexpressing muscle (Fig. 4E, F), and that cleavage of the full-length GFP-mCherry-Atg8a that occurs in the lysosome was inhibited (Fig. 4G). These data all indicate that of maturation of autophagosomes to autolysosomes is impaired. Finally, complementing these observations with organism-scale genetic manipulations, we found that overexpression of *p62* or *Atg8a* was sufficient to significantly rescue the climbing deficit caused by *Cisd* overexpression (Fig. 4H).

Taken together, these data suggest that the Cisd-related block in mitophagy is caused by an impairment in autophagosome engulfment of defective mitochondria, which ultimately results in autophagosome accumulation and a blockage of general autophagy flux.

#### Reducing Cisd levels promotes mitophagy in an age and context-dependent manner

Since Cisd accumulation blocks mitophagy, we surmised that reducing Cisd levels could have the opposite effect and facilitate mitophagy. To approach this, we first assessed available loss-of-function reagents for *Cisd*. A null mutant, *Cisd^−/−^*, had significant impact on organismal and mitochondrial phenotypes, conferring shorter lifespan, reduced climbing ability and fragmented mitochondria (Fig. S2A-D), consistent with a previous report [36]. Two independent transgenic *Cisd* RNAi constructs (termed KK and GD), which substantially reduced the Cisd protein levels (Fig. S2A), produced similar but generally milder phenotypes (Fig. S2B-D). Hence, for reasons of greater versatility (i.e., tissue-specific targeting) the RNAi lines were used for subsequent studies.

Analysing mitophagic flux using the *mito*-QC reporter, we observed a significant increase in mitophagy with age both in muscle (Fig. 5A, B) and neurons (Fig. S2E, F). Interestingly, *Cisd* knockdown was able to significantly increase mitophagy specifically in muscle tissue of older flies (Fig. 5A, B). No induction was observed in young flies or in aged neurons (Fig. S2E, F).

**Figure 5.**
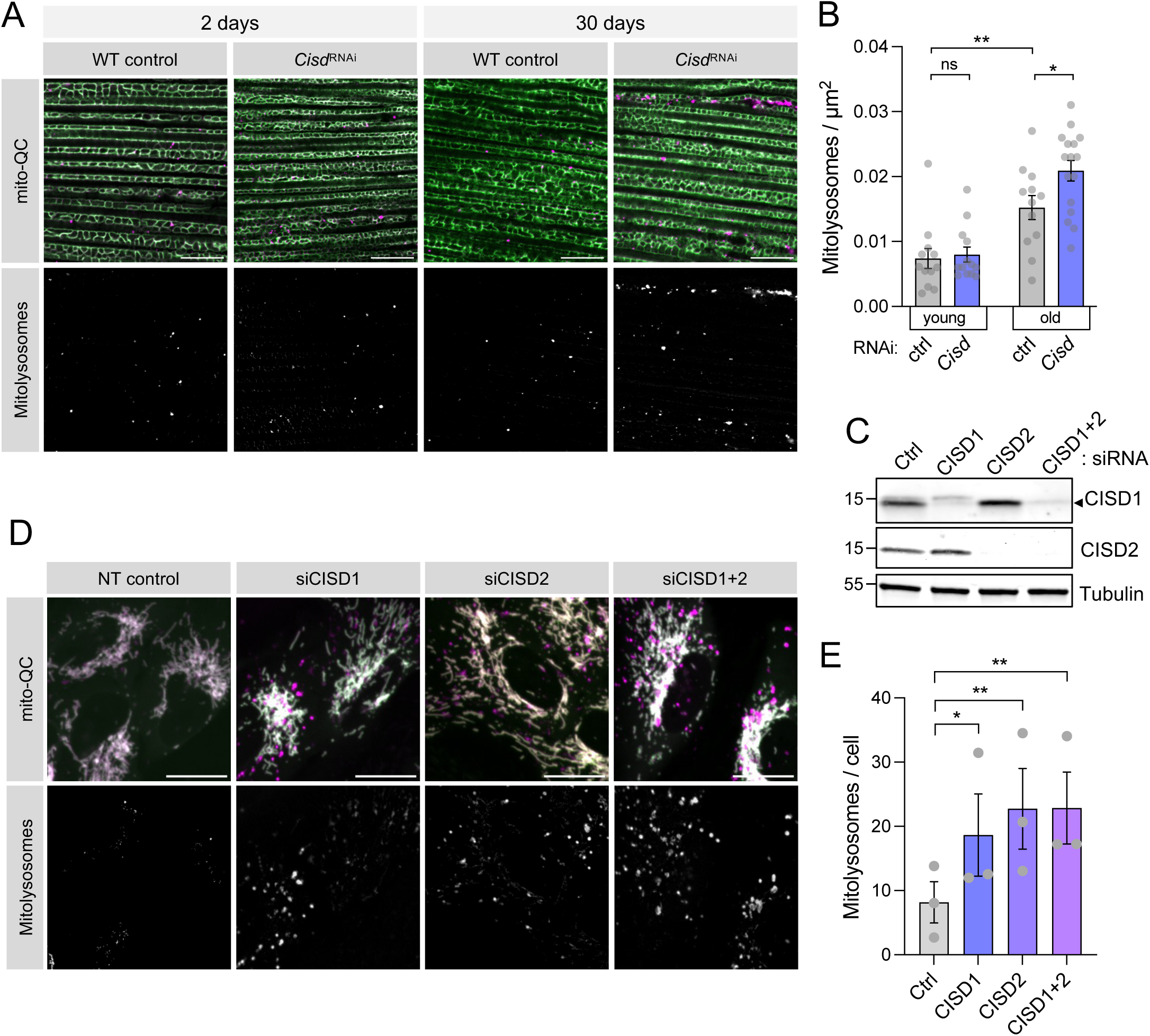
Loss of *Cisd* promotes mitophagy flux. (A, B) Confocal microscopy analysis of mitophagy reporter *mito*-QC in flight muscle from WT control and *Cisd* knockdown flies of the indicated ages. (B) Quantification of the number of mitolysosomes shown in A. Data points indicated individual animals analysed. Statistical analysis: unpaired t-test; * *P* < 0.05; ** *P* < 0.01. (C) Immunoblot analysis of ARPE-19 cells expressing *mito*-QC, shown in D, with non-targeting siRNAs (Ctrl) or targeting CISD1, CISD2 or both. (D) Confocal microscopy analysis of mitophagy using *mito*-QC in cells shown in C. (E) Quantification of the number of mitolysosomes shown in D. Data points indicated replicate experiments. Statistical analysis: one-way ANOVA with Dunnett’s post-hoc correction; * *P* < 0.05; ** *P* < 0.01. Scale bars = 10 μm.

Seeking to validate these results in human cells, we knocked down CISD1 and/or CISD2 in a mitophagy reporter cell line (ARPE-19-*mito*-QC) (Fig. 5C) and assessed the formation of mitolysosomes. In agreement with the *in vivo* data, we found that siRNA-mediated depletion of CISD1, CISD2 or both fragmented mitochondria and significantly increased basal mitophagy (Fig. 5D, E). Together, these data indicate that reducing CISD levels can increase mitophagy in *Drosophila* and human cells.

### Reducing Cisd suppresses *Pink1/parkin* mutant phenotypes by upregulating mitophagy

We have established that Cisd is a key Parkin substrate that accumulates in *Pink1/parkin* mutants and its pathologic build-up induces mitochondrial disruption and neurodegenerative phenotypes. Therefore, we reasoned that reducing Cisd levels in *Pink1/parkin* mutants could ameliorate the pathological phenotypes. *Pink1/parkin* mutants exhibit an array of well characterised neurodegenerative phenotypes, including locomotor deficits, degeneration of flight muscle accompanied by gross mitochondrial disruption, progressive dopaminergic (DA) neuron loss, and shortened lifespan [13–15]. Ubiquitous knockdown of *Cisd* in *parkin* or *Pink1* mutants significantly suppressed their climbing defects (Fig. 6A, B) and disruption of flight muscle mitochondria (Fig. 6C, D). The two RNAi lines suppressed equally well, so for simplicity subsequent experiments were conducted with the KK RNAi line. *Cisd* knockdown also significantly suppressed the degeneration of DA neurons (Fig. 6E, F) and shortened lifespan (Fig. 6G, H) in both *parkin* and *Pink1* mutants. Interestingly, muscle-specific depletion of *Cisd* was sufficient to partially rescue the locomotor phenotype of *parkin* mutants while pan-neuronal knockdown did not (Fig. S3A, B). Overall, these data demonstrate that reduction in Cisd levels is sufficient to significantly reduce the pathology associated with loss of *Pink1* or *parkin*. Supporting the genetic interaction between *Cisd* and *Pink1/parkin*, it was notable that *Cisd* overexpression, while viable in a wild-type background, caused complete lethality in *Pink1/parkin* mutants (Fig. S3C). Thus, there exists a strong genetic interaction between *Cisd* and *Pink1/parkin* implicating a strong functional interaction.

**Figure 6.**
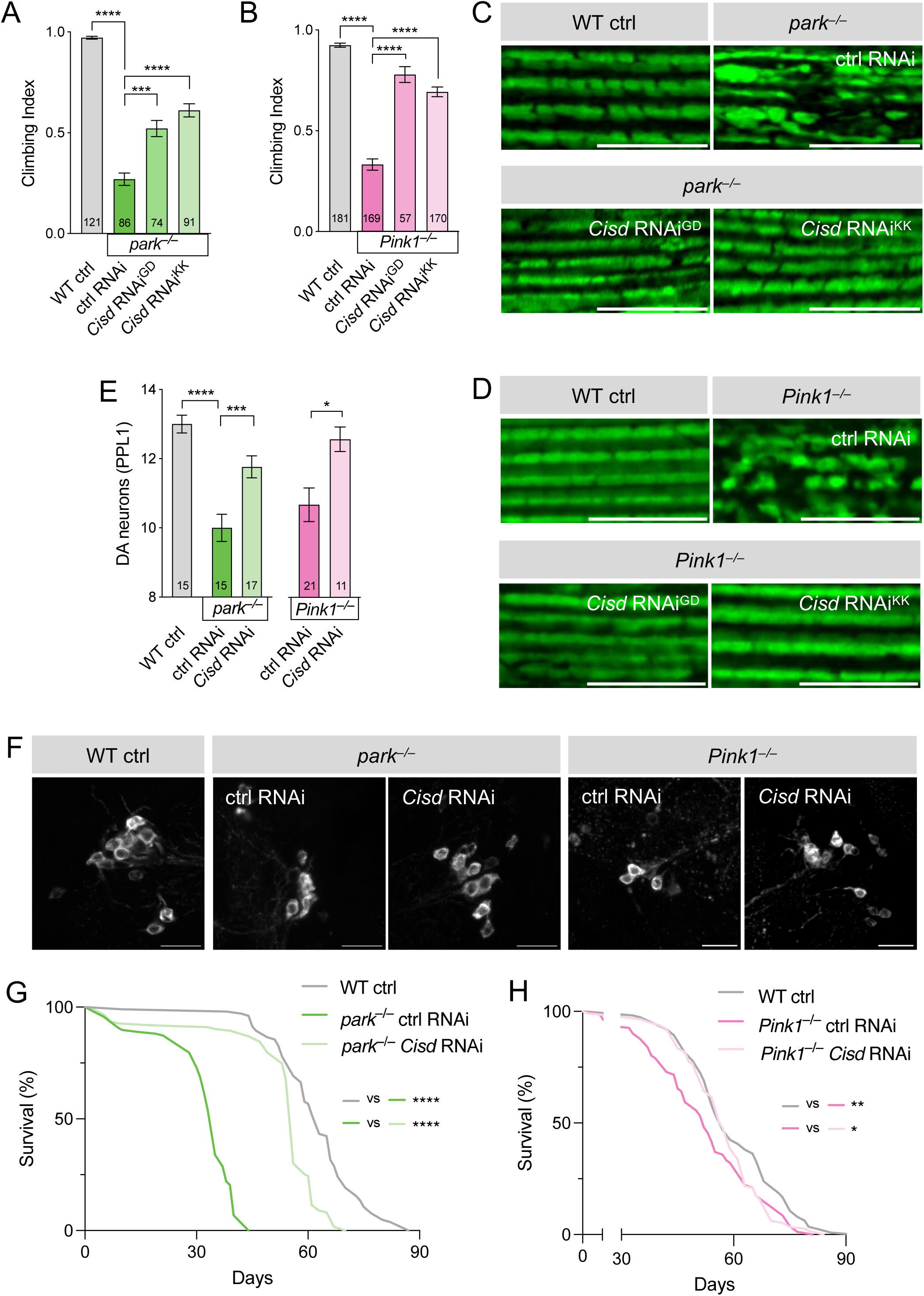
*Cisd* knockdown ameliorates *Pink1/parkin* mutant phenotypes. (A, B) Climbing analysis of WT control versus (A) *parkin* or (B) *Pink1* mutants with control or *Cisd* RNAi. GD and KK indicate independent RNAi lines. Charts show mean ± 95% CI. N is shown inside bars. (C, D) Confocal microscopy of flight muscle from (young) wild-type control (WT ctrl) or (C) *parkin* or (D) *Pink1* mutants co-expressing with control or *Cisd* RNAi and the mito.GFP mitochondrial marker. (E) Quantification of dopaminergic (DA) neurons shown in F. F) 30-day-old WT or *parkin* or *Pink1* mutants co-expressing control or *Cisd* RNAi, immunostained for tyrosine hydroxylase. (G, H) Lifespan analysis of WT versus (G) *parkin* or (H) *Pink1* mutants with control or *Cisd* RNAi. Statistical analyses: (A, B) Kruskal-Wallis non-parametric test with Dunn’s post-hoc correction, (E) one-way ANOVA with Sidak’s post-hoc correction; (G, H) Log rank (Mantel-Cox) test. * *P* < 0.05; ** *P* < 0.01; *** *P* < 0.001; **** *P* < 0.0001. Scale bars = 10 μm.

Having shown that loss of *Cisd* promotes mitophagy in wild-type flies and human cells, we assessed whether reducing Cisd increased mitophagy in the *Pink1/parkin* mutants. As expected, we found that mitochondria from *Pink1* and *parkin* mutants accumulate high levels of p62 and Atg8a-II (Fig. 7A, B), consistent with a block in mitochondrial turnover, as also occurs upon *Cisd* overexpression (Fig. 4D). In this context, *Cisd* knockdown prevented the aberrant build-up of p62 and Atg8a-II in the *Pink1/parkin* mutants (Fig. 7A, B). Moreover, *Cisd* knockdown also substantially reduced the dramatic build-up of pUb that occurs in *parkin* mutants (Fig. 7C). Immunostaining thoraces of these flies showed similar depositions of p62 aggregates *Pink1* and *parkin* mutants, which were also substantially reduced upon *Cisd* knockdown (Fig. 7D). The reduction in p62 build-up correlated with a notable improvement in mitochondrial morphology, labelled with ATP5a (Fig. 7D). Investigating mitophagy flux directly using the *mito*-QC reporter, we found that indeed reducing *Cisd* increased mitophagy in *parkin* mutant muscle (Fig. 7E, F), which likewise correlated with improved mitochondrial morphology and general tissue health.

**Figure 7.**
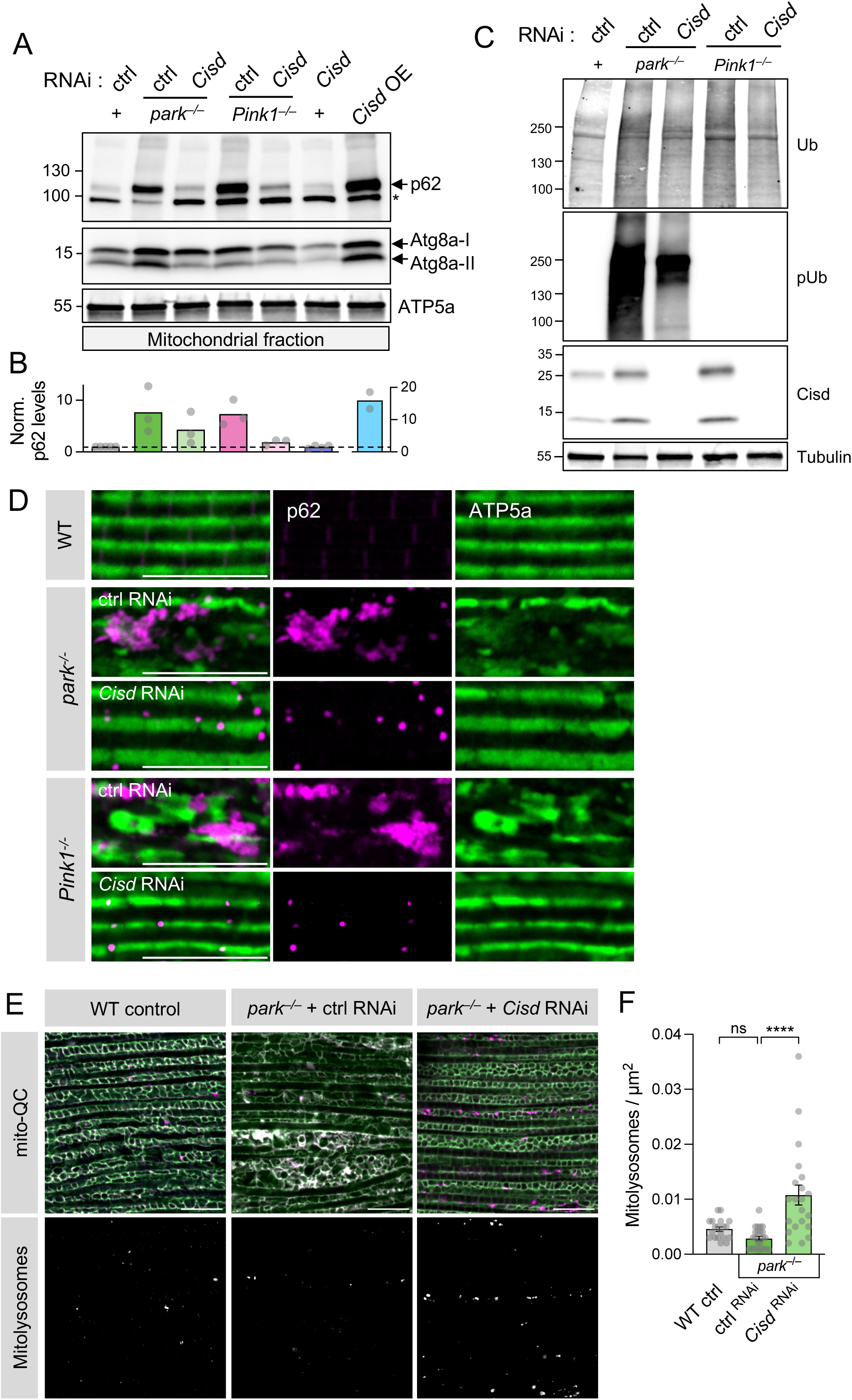
Loss of *Cisd* rescues *Pink1* and *parkin* mutant degenerative phenotypes. (A) Immunoblot of mitochondrial protein lysates from *Cisd* knockdown in WT versus *Pink1* and *parkin* mutant backgrounds, alongside respective controls, probed for p62 and Atg8a (LC3) levels with ATP5a loading control. (B) Relative p62 levels quantified from blots in A, normalised to WT control. (C) Immunoblots of whole fly lysates of genotypes as in A, probed for pUb and respective controls. (D) Confocal micrographs of adult flight muscle *Cisd* knockdown in WT versus *Pink1* and *parkin* mutant backgrounds, alongside respective controls, immunostained for APT5a and p62. (E, F) Confocal microscopy analysis of mitophagy reporter *mito*-QC in flight muscle from *Cisd* knockdown in WT and *parkin* mutant backgrounds, alongside WT control. (F) Quantification of the number of mitolysosomes shown in E. Data points indicated individual animals analysed. Statistical analysis: one-way ANOVA with Sidak’s post-hoc correction; **** *P* < 0.0001. Scale bars = 10 μm.

Taken together, these data demonstrate that Cisd-depletion induces upregulation of mitophagy and rescues the abnormal accumulation of pUb, p62 and Atg8a back to basal levels, preventing neurodegenerative phenotypes in *Pink1* and *parkin* mutant flies.

### The CISD inhibitor rosiglitazone induces mitophagy and is beneficial for *Pink1/parkin* **mutant phenotypes**

Given that genetic reduction of *Cisd* levels robustly rescued *Pink1/parkin* phenotypes, we considered the potential for putative CISD1/2 small-molecule inhibitors to have similar beneficial effects. Several compounds, including rosiglitazone, pioglitazone and NL1, have been reported to potently inhibit CISD1 and CISD2 [43]. Following our findings that CISD1/2 knockdown was sufficient to induce mitophagy, we tested the effects of two structurally distinct compounds, rosiglitazone and NL1, for their potential to induce mitophagy. Exposing mitophagy reporter cells to rosiglitazone and NL1 for 24h, we found that both compounds intensely induced mitophagy as well as mitochondrial fragmentation (Fig. 8A, B), mirroring the effects of genetically reducing *CISD1*/2.

**Figure 8.**
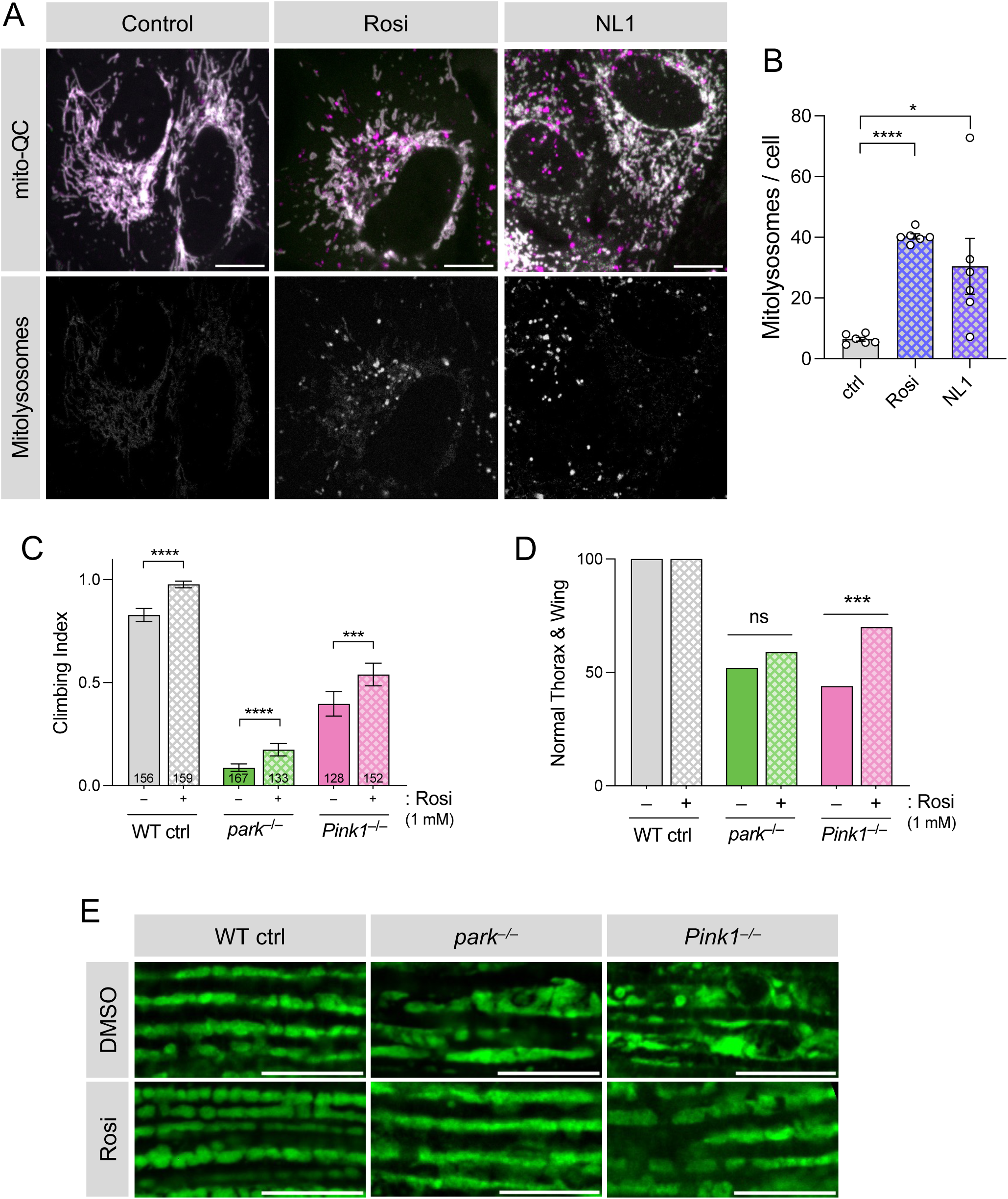
CISD inhibitors induce mitophagy and rescue rescues *Pink1* and *parkin* mutant phenotypes. (A) Confocal microscopy analysis of WT ARPE-19 cells expressing *mito*-QC to visualise mitolysosomes (shown separately) treated with 100 µM rosiglitazone (Rosi), NL1 or vehicle. (B) Quantification of the number of mitolysosomes per cell of conditions shown in A. Data points indicated replicate experiments. Statistical analysis: RM one-way ANOVA with Geisser-Greenhouse correction; * *P* < 0.05; **** *P* < 0.0001. (C) Analysis of Climbing, (D) thoracic indentations and drooped-wing phenotype, and (E) mitochondrial morphology in flight muscle of *Pink1* and *parkin* mutants alongside WT control flies, treated with 1 mM rosiglitazone (Rosi) or vehicle. Statistical analysis: Chi-squared test. *** *P* < 0.001; **** *P* < 0.0001. Scale bars = 10 μm.

These results motivated us to test the therapeutic potential of CISD inhibitors *in vivo*. The preceding results showed that rosiglitazone had an overall greater and more consistent effect on mitophagy induction than NL1, hence, we focussed on rosiglitazone for *in vivo* analysis. To this end, we treated *Pink1* and *parkin* mutants with rosiglitazone-dosed food and examined their classic phenotypes. Importantly, *Pink1/parkin* mutants treated with rosiglitazone showed a modest but significant improvement in their climbing ability (Fig. 8C). There was also a general modest improvement of thoracic indentations and abnormal wing posture (Fig. 8D), caused by the degeneration of underlying flight muscle, rosiglitazone treatment visibly improved mitochondrial morphology of *Pink1/parkin* flight muscle (Fig. 8E). Together, these results support the potential benefit of CISD inhibitors in promoting mitophagy and alleviating degenerative phenotypes in these PD models.

## Discussion

In this study, we have shown that *Drosophila* Cisd accumulates in *Pink1/parkin* KO flies, as might be expected from a degradation target, but it also accumulates during normal wild-type ageing. We have demonstrated that *Cisd* overexpression is sufficient to dramatically affect mitochondrial morphology, motor behaviour and lifespan, and is particularly toxic when accumulating in neurons. Considering the conservation with mammalian CISD1/2, our results indicate that *Drosophila* Cisd functions like human CISD1 rather than CISD2, and overexpressing human CISD1 similarly impacts mitochondrial morphology *in vivo*. Mechanistically, we have found that *Cisd* overexpression blocks mitophagy and inhibits autophagic flux, and *in vivo* overexpression phenotypes can be suppressed by upregulation of key autophagy regulators. Importantly, we showed that loss of CISD1/Cisd increased mitophagy in human cells and *in vivo*. Consistent with these effects, genetic loss of *Cisd* significantly rescued *Pink1/parkin* mutant phenotypes, including mitochondrial integrity and DA neurodegeneration. Finally, we demonstrated that known small-molecule CISD inhibitors are able to induce mitophagy in cells and significantly rescue *Pink1/parkin* fly phenotypes, mirroring the genetic reduction. Hence, we posit that CISD1/Cisd proteins play an important role in regulating mitochondrial function and quality control, ultimately impacting on age-related neurodegeneration.

The [2Fe-2S] cluster-coordinating CISD proteins act as multi-functional regulators of important cellular processes including redox balance, iron metabolism, calcium signalling and mitochondrial respiration [25]. One of the most striking recent developments in understanding the role of CISD proteins is their impact on mitochondrial morphology and dynamics [26,29,36,44–46]. In general, loss of CISD proteins causes mitochondrial fragmentation while their accumulation causes clumping and hyperfusion. Mitochondrial dynamics plays an intimate in role in regulating mitochondrial quality control and neuronal survival. Notably, mitochondrial fragmentation is known to facilitate mitophagy [47], consistent with our observations of *CISD1/Cisd* reduction. How CISD proteins mediate effects on mitochondrial morphology is not clear, however, CISD1 has been shown to promote inter-mitochondrial junctions (IMJs) [44], where cristae of neighbouring mitochondria align, likely as a way to couple and enhance respiration [48]. We hypothesise that the enlarged/clumped mitochondrial morphology we observe upon *CISD1/Cisd* overexpression may be a consequence of increased IMJs induced by the strong Cisd dimerisation.

Indeed, our finding that Cisd protein levels increased with age, also noted elsewhere [36], suggests further mechanistic links between CISD proteins and age-related mitochondrial disruption. The reason behind the accumulation is unclear at present but could reflect a decline in parkin activity with age or a compensatory response for an increased need for Cisd activity. Nevertheless, since multiple studies have shown that mitochondria become enlarged with age [36,49–52], and some have linked this to defective autophagy and/or Parkin function [13–15,41,46,50,52], it is tempting to speculate that the accumulation of CISD proteins may be a key factor driving mitochondrial clumping/enlargement during ageing. Alternatively, consistent with our data showing that Cisd accumulation inhibits autophagic flux, others report that CISD1/2 may directly regulate autophagy, possibly via modulating Ca^2+^ signalling [33,35,46,52]. Since evidence indicates that proteostasis and autophagy decline with age [53], it is also possible that ageing related decrease in autophagic capacity could also be directly related to increased CISD protein levels.

The abundant evidence that mitochondrial disruption contributes to multiple neurodegenerative diseases [54] and ageing itself [53] has prompted intense interest in identifying ways to upregulate mitophagy as a therapeutic intervention. We have shown that depletion of CISD proteins increased mitophagy *in vivo* and in human cells. A recent report has also described that genetic or pharmacologic inhibition of CISD1 upregulated Parkin-dependent mitophagy in human cells [55], although our data indicate that *Cisd* depletion-induced mitophagy can be Parkin-independent since it occurred in *parkin* mutant flies. Interestingly, while we found that *Cisd* depletion-induced mitophagy alleviated blocked mitochondrial turnover in *Pink1/parkin* mutants, under steady-state conditions mitophagy induction occurred in quite selective circumstances, notably in aged muscle. Surprisingly, *Cisd* depletion did not have a significant effect on neuronal mitophagy. It is becoming clear that mitophagy rates can greatly vary between tissues [8,9,11,56], and the neurons analysed here displayed higher levels of basal mitophagy (even in young individuals) compared to muscle. Indeed, the differing mitophagy induction perfectly correlated with *Cisd* RNAi being able to rescue *parkin* mutant locomotion defects when expressed in muscle but not in neurons. Altogether, these data suggest that *Cisd* depletion-mediated mitophagy likely occurs in conjunction with additional stress conditions, for example, with ageing or *Pink1/parkin* related pathology. It will be important for future work to unravel the molecular mechanism of how CISD proteins regulate mitophagy and the circumstance in which this can be unleashed for therapeutic benefit.

Importantly, CISD proteins are known to be targeted by small-molecules currently used as anti-diabetic drugs. CISD1 was originally identified as a target of the thiazolidinedione (TZD) drug, pioglitazone [57], and subsequent studies showed that several distinct TZDs target CISD1 and CISD2 [43]. We have shown that pharmacologic inhibition of CISD proteins via anti-diabetic drugs promotes mitophagy in human cells and ameliorates *Pink1/parkin* mutant phenotypes. While additional work is required to fully evaluate the therapeutic potential of these compounds, it is intriguing to note that epidemiological studies have linked diabetes with an increased risk of PD [58], and diabetic patients treated with anti-diabetic drugs have a reduced risk of developing PD [59,60].

## Conclusion

Cisd naturally accumulates in *Drosophila* tissues during aging and in *Pink1/parkin*-mutant PD model flies, causing mitochondrial defects that result in mitophagy and autophagy impairment. Genetically or pharmacologically reducing Cisd activity, alleviates age-related neurodegenerative pathology by upregulating mitophagy, a phenomenon which is conserved in human cells. Thus, we propose that inhibiting CISD proteins, for which FDA-approved drugs are already available, represents a potential therapeutic target for the treatment of neurodegenerative diseases, such as PD but also for age-related decline. Future work is needed to better define the molecular mechanism by which mitophagy is upregulated upon CISD1/2 inhibition and to refine the pharmacokinetics and specificity of potential new inhibitors.

## Availability of data and materials

All data needed to evaluate the conclusions in the paper are present in the paper and/or the Supplementary Materials. This study includes no data deposited in external repositories. Additional data related to this paper may be requested from the authors.

### Abbreviations

(BSA): Bovine Serum Albumin
(DA): Dopaminergic
(FBS): Fetal Bovine Serum
(FA): Formaldehyde
(MQC): Mitochondrial quality control
(OMM): Outer mitochondrial membrane
(PD): Parkinson’s Disease
(PBS): Phosphate buffered saline
(pUb): Ser65-phospho-Ubiquitin
(TZD): Thiazolidinedione
(WT): Wild-type

## Acknowledgements

We kindly thank Dr Ian Ganley (MRC Protein Phosphorylation and Ubiquitylation Unit, University of Dundee) for proving us with the ARPE-19-mito-QC cells. We would also like to thank Prof. Jon Lane (University of Bristol) for generously donating us the hTERT-RPE1 and hTERT-RPE1-YFP-PARKIN cells. Jan Miljkovic, Jordan Morris, especially Michele Frison for valuable input, and all the members of the Whitworth’s lab for discussions.

## Contributions

AM and AJW conceived the project; AM, ASM and AW designed the experiments; AM, ASM, JTP, MJT, ATF and PLC performed experimental work; CHC provided key advice, input and reagents; AM, ASM and AW analysed data; AM and AJW wrote the manuscript with input of all co-authors; AJW supervised the project. All authors read and approved the final manuscript.

### Corresponding author

Correspondence to <u>Aitor Martinez</u> and <u>Alex Whitworth</u>.

### Ethics declarations

#### Ethics approval and consent to participate

Not applicable.

#### Consent for publication

Not applicable.

---

**Supplementary Figure S1.**
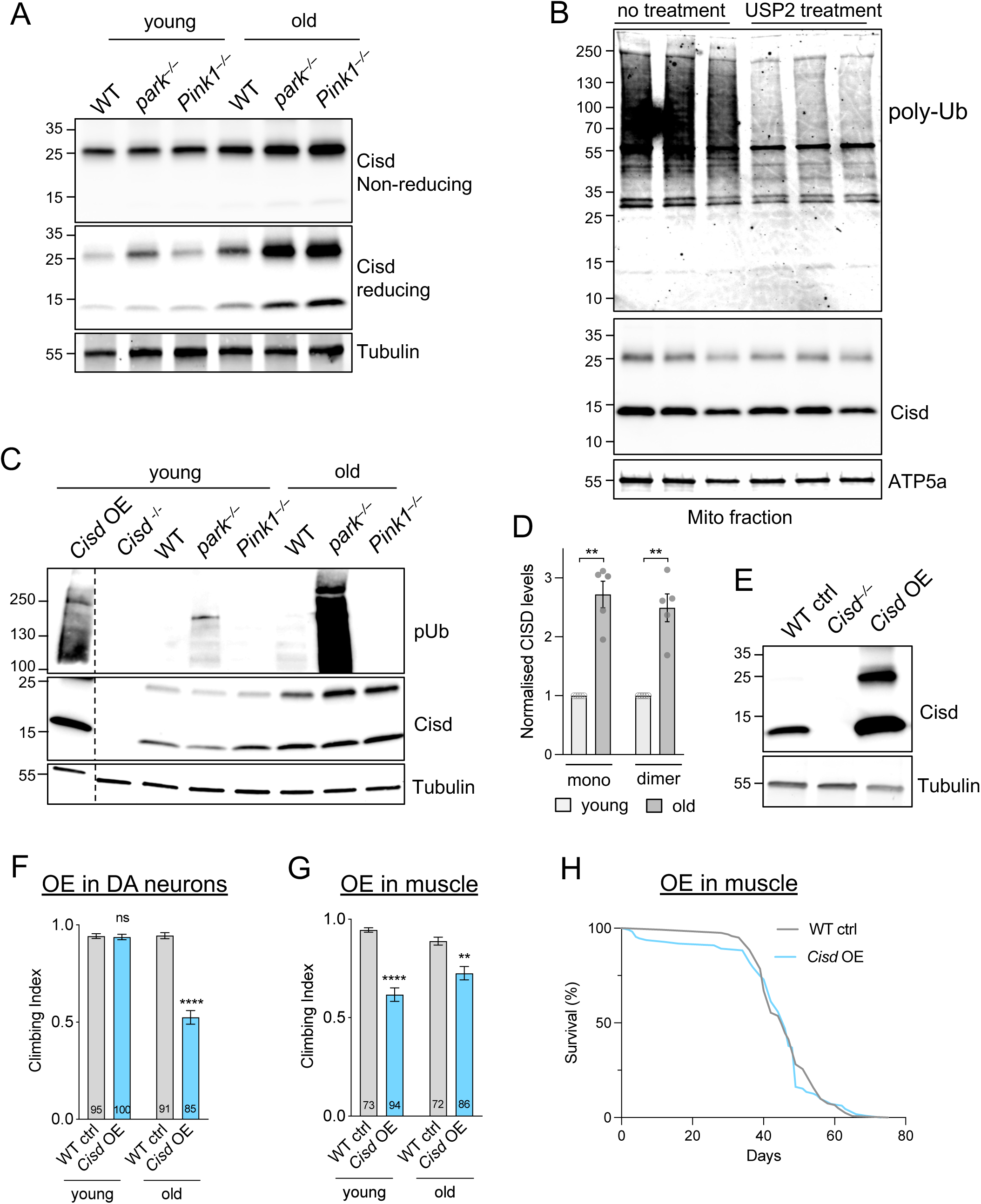
Characterisation of *Cisd* in ageing and different tissues. (A) Immunoblot analysis of protein lysates from young (2-day-old) or old (30-day-old) whole flies of the indicated genotypes. Samples were homogenised under non-reducing (top panel) and reducing (bottom panel) conditions. Blots were probed for Cisd and Tubulin. (B) Immunoblot of mitochondrial fractions from WT flies, probed for poly-Ub, Cisd and ATP5a. (C) Immunoblot analysis of protein lysates from young (2-day-old) or old (30-day-old) whole flies of the indicated genotypes, probed for pUb, Cisd and Tubulin. (D) Relative amount of Cisd monomer and dimer quantified from replicate blots shown in C. (E) Immunoblot of protein lysates from whole flies of WT or *Cisd* null (−/−) or *Cisd* overexpression (OE), probed for Cisd and Tubulin. (F, G) Climbing assay of young and old WT flies or *Cisd* overexpression driven only in (F) DA neurons (*TH*-GAL4) or (G) pan-muscle (*Mef2*-GAL4). (H) Lifespan analysis of flies overexpressing *Cisd* in muscles as in G versus a WT control genotype. Statistical analysis: (D) unpaired t-test; (F, G) Kruskal-Wallis non-parametric test with Dunn’s post-hoc correction. ** *P* < 0.01, **** *P*<0.0001.

**Supplementary Figure S2.**
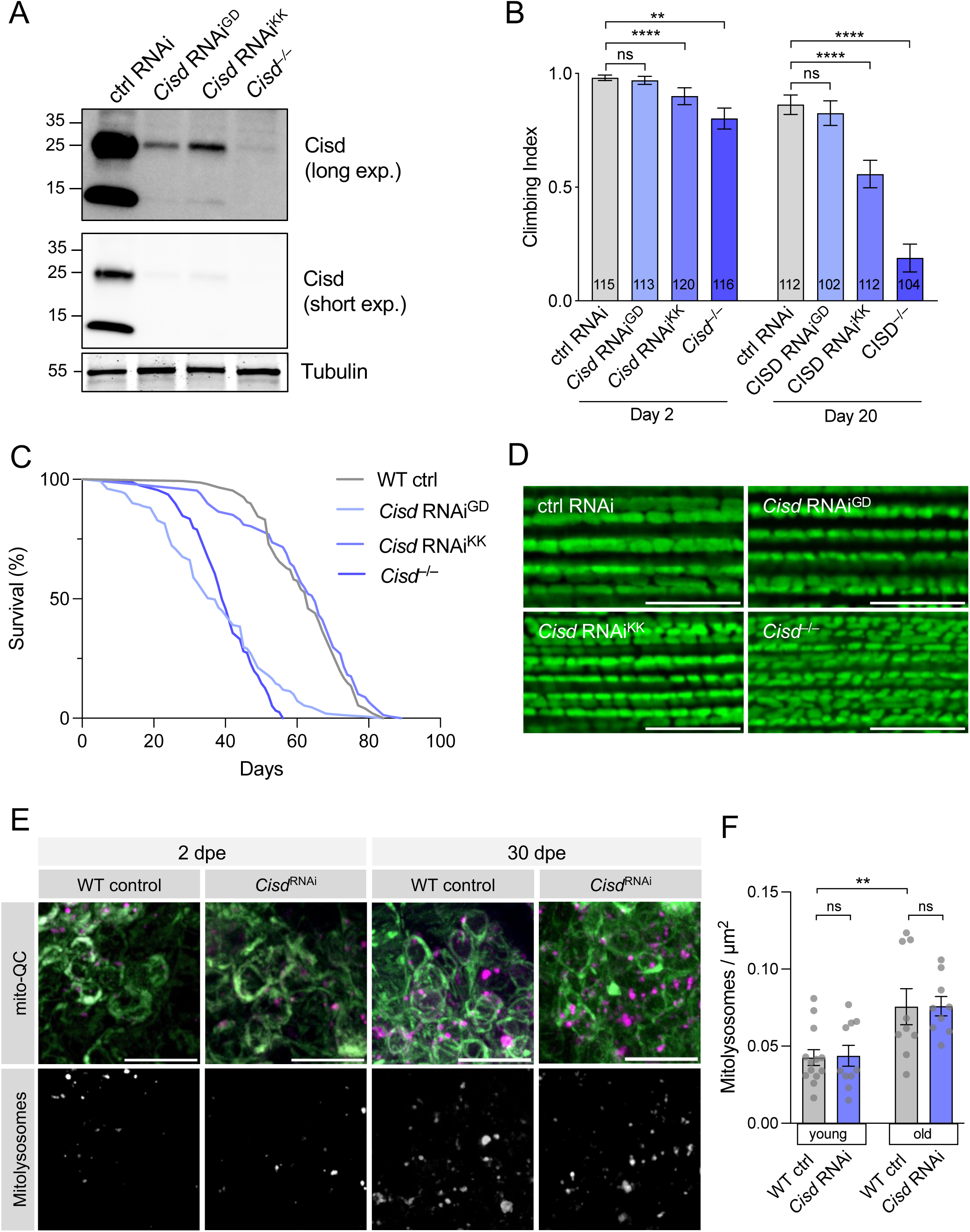
Characterisation of *Cisd* loss-of-function on organismal and mitochondrial phenotypes. (A) Immunoblot analysis of protein lysates from whole flies of the indicated genotypes of *Cisd* loss versus control, probed for Cisd and Tubulin. (B) Climbing assay of young (2-day-old) or older (20-day-old) flies of the indicated genotypes. (C) Lifespan analysis of *Cisd* loss as in B. (D) Confocal microscopy analysis of mitochondrial morphology, immunostained for ATP5A mitochondrial marker in flight muscle of the indicated genotypes. (E) Confocal analysis of adult neurons of the indicated ages, WT control and *Cisd* RNAi animals co-expressing the mitophagy reporter *mito*-QC (OMM-localised tandem RFP-GFP) to highlight mitolysosomes, shown separately and quantified in F. Statistical analysis: (B) Kruskal-Wallis non-parametric test with Dunn’s post-hoc correction. (F) one-way ANOVA with Sidak’s post-hoc correction. ** *P* < 0.01, **** *P*<0.0001. Scale bars = 10 μm.

**Supplementary Figure S3.**
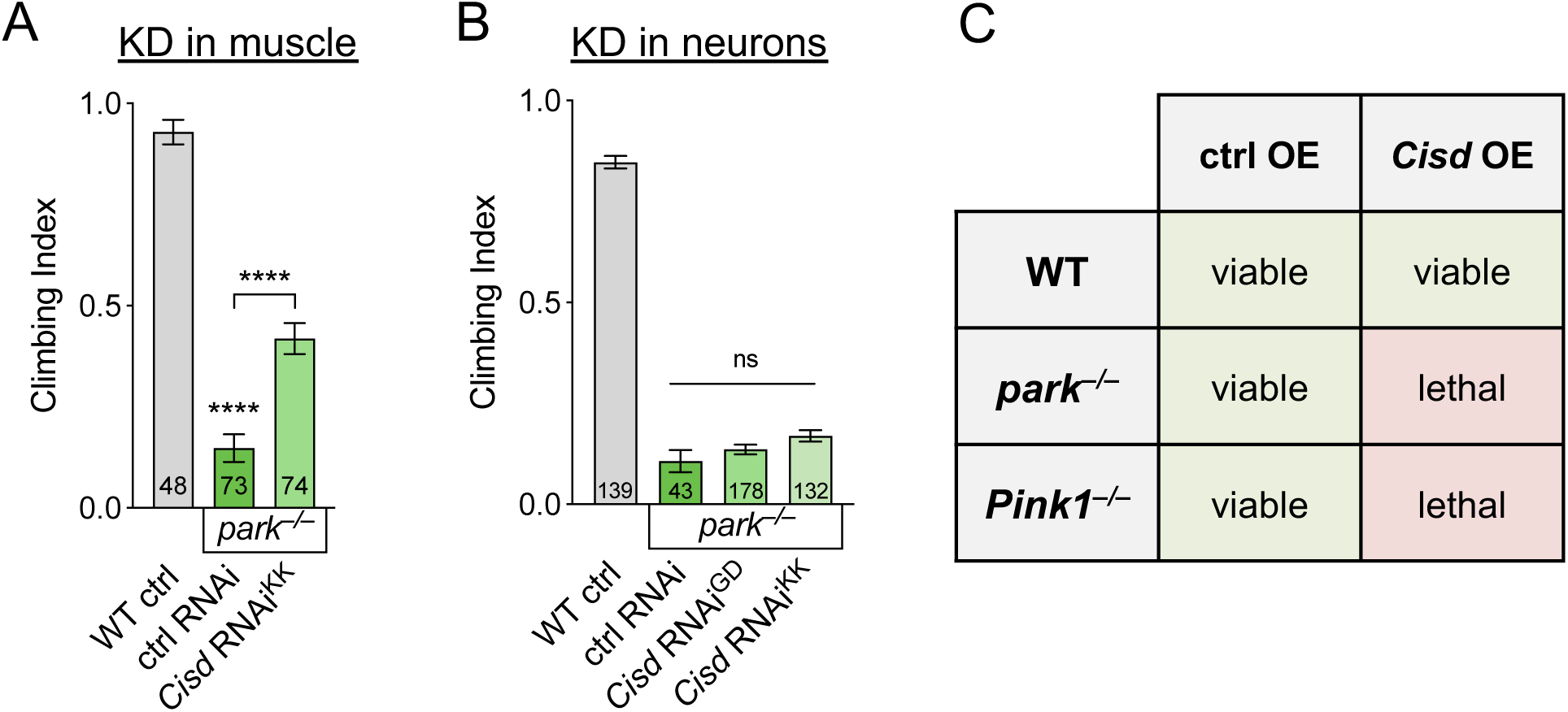
Characterisation of *Cisd* and *parkin* genetic interaction. (A, B) Climbing assay of tissue-specific *Cisd* knockdown (KD) in either pan-muscles (A) or pan-neurons (B) alongside respective controls. (C) Viability assay for genetic interactions between the indicated combinations revealing synthetic lethality of *Pink1/parkin* mutants and *Cisd* OE.

